# Characterization of heterogeneous bioinformatics training needs in the AI era

**DOI:** 10.64898/2026.10.22.761024

**Authors:** Geert van Geest, Valeria Di Cola, Monique Zahn, Diana Marek, Gregoire Rossier, Wandrille Duchemin, Joana Carlevaro-Fita, Deepak Tanwar, Marion Vaucher, Patricia M. Palagi

**Affiliations:** Training Group, SIB Swiss Institute of Bioinformatics, Amphipôle, Quartier UNIL-Sorge, 1015 Lausanne, Switzerland; Swiss Biobanking, Biopôle – Bâtiment Phényl, Route de la Corniche 3A, 1066 Epalinges, Switzerland; Vital-IT Group, SIB Swiss Institute of Bioinformatics, Amphipôle, Quartier UNIL-Sorge, 1015 Lausanne, Switzerland; sciCORE Center for Scientific Computing, University of Basel, Klingelbergstrasse 61, 4056 Basel, Switzerland; Interfaculty Bioinformatics Unit, University of Bern, Baltzerstrasse 6, 3012 Bern, Switzerland; Department of Quantitative Biomedicine, University of Zurich, Winterthurerstrasse 190, 8057 Zurich, Switzerland

**Keywords:** training need assessment, learner profiles, network analysis, cluster analysis, curriculum design

## Abstract

Bioinformatics training providers serve heterogeneous audiences, and generative artificial intelligence (AI) is rapidly changing what learners need. Training-needs surveys are typically analyzed question by question or split by demographics such as career stage. This assumes that people in the same demographic group need the same training, whereas needs may depend more on the data people work with. We present a protocol that groups survey respondents by the similarity of their topic ratings rather than by demographics. Respondents form a correlation network, and community detection identifies learner profiles with shared needs. Applied to the 2026 SIB Swiss Institute of Bioinformatics training-needs survey (n=260), it yielded four profiles that were stable under resampling: AI and data, infectious disease genomics, functional genomics, and biochemistry, imaging & AI, each with distinct preferences and barriers. For example, the topic ‘LLM-based knowledge extraction’ ranked 2nd overall but 16th for the infectious disease genomics profile, where ‘phylogenetics data analysis’ (17th overall) ranked first. Across all profiles, respondents wanted training on AI and reproducible data practices but liked AI-assisted learning least of all formats. The protocol requires only Likert-type items on a shared scale, so other providers can reuse it to shape their training curricula.

**Biographical note:** the authors are members of the SIB Training group that provides training to ensure that life scientists make the best of the latest advances in computational biology.

**Key points:**

1. Training-needs surveys are typically analyzed question by question or by demographic group, which can hide the distinct needs of subgroups within heterogeneous bioinformatics audiences.
2. We present a network-clustering method for identifying learner profiles based on survey data.
3. Applied to the 2026 SIB training-needs survey, the method identified four thematically distinct, reproducible learner profiles, capturing training-need heterogeneity that demographics miss.
4. AI/LLM skills are the top training gap across all profiles, alongside strong demand for FAIR/reproducible data practices — yet respondents prefer human-led formats over AI-assisted learning.
5. Curricula should be tailored to learner profiles rather than demographics or aggregate topic rankings, which each hide profile-specific needs.

## 1 Introduction

The rapid and continuous evolution of high-throughput experimental technologies, large-scale data infrastructures, and artificial intelligence (AI) makes bioinformatics training a necessity for life science researchers [1]. National and international training providers, such as the SIB Swiss Institute of Bioinformatics (SIB), aim to provide high-quality, effective short-format courses [2] to a heterogeneous audience at point of need. This audience ranges from academic PhD candidates requiring upskilling in specific analytical competences to corporate senior scientists or clinical personnel seeking broad overviews of emerging resources in computational biology.

It is a continuous challenge to match the training needs of such a diverse audience. Training-needs surveys are frequently used to understand these [e.g. 3,4–8]. These previous training-needs surveys were analyzed using various dimensions such as course topics, format, and logistics independently or using classical demographic categories like career stage or geographic region to slice responses. While informative, such an approach assumes that individuals within a demographic sub-group are homogeneous. However, demographic characteristics like career stage, region or sector do not necessarily explain common training needs; a post-doctoral researcher likely shares more training needs with a research technician in industry who works with similar data, than with a post-doctoral researcher working in a different domain.

Clustering of survey responses can be highly valuable for understanding such respondent heterogeneity. The approach was already introduced in the eighties for market segmentation [9] and currently finds a wide range of applications [10–14]. For our case of bioinformatics training needs, identifying clusters can reveal survey respondents with similar needs. Therefore, it can help training providers design curricula that are tailored to the specific needs of these sub-groups, complementing information obtained from broad demographic data.

In recent years, the fast growth of AI and LLMs has transformed both the bioinformatics landscape and the nature of computational training itself [15,16]. Various generative AI tools have shifted how researchers write code [17], automate workflows and retrieve data [18]. LLMs and generative AI are heavily used by scientists [19], and they are frequently used for education [20,21], both in the classroom and for self-directed learning [22,23]. Identifying which AI-related topics scientists actually need would let training providers design courses around those needs and integrate AI topics across the curriculum rather than confining them to dedicated AI courses.

This paper describes a data-driven framework for understanding training needs based on the SIB training-needs survey in terms of course topics, formats, and barriers. By converting individual responses into a correlation graph, we identify clusters of respondents who share common training needs, regardless of their institutional or regional backgrounds, and characterize each cluster as a learner profile. We detail the underlying data preprocessing pipeline, a conceptual overview of graph construction, and the subsequent characterization of these learner profiles, with an emphasis on how AI can be incorporated in training programs.

## 2 Methods

### 2.1 Survey design and distribution

The SIB training-needs survey was designed to collect information using mostly multiple-choice and Likert-type questions. All survey questions, and anonymized responses, can be found in Supplementary Data 1. At its core, the questionnaire evaluated course topics, barriers and formats using 5-point Likert-type scales:

- Training needs: Respondents evaluated their training needs for 33 topics nested within five global computational domains: *Omics analysis*, *Computational methods and AI*, *Data management and knowledge representation*, *Biomedicine and pathogens*, and *Biodiversity and Ecology*, which were quantified using an ordinal scale ranging from ‘Not needed’ to ‘Absolutely essential’.
- Perceived participation barriers: Participation barriers like course fees, travel distance, timing constraints, and prerequisite alignment were quantified using an ordinal scale ranging from ‘No barrier’ to ‘Critical barrier’.
- Preferred learning formats: Delivery formats like webinars, bring your own data, problem-based learning, multi-day intensive courses, self-paced e-learning, and AI-assisted learning were quantified using an ordinal scale ranging from ‘Strongly dislike’ to ‘Strongly like’.

Demographic data were also collected through multiple-choice questions to characterize the respondent pool: career stage, geographic demographics (working country and Swiss canton), organization type (Academia, Healthcare, Industry, Non-profit), and historical interaction (time elapsed since attending their last SIB training course).

The survey was created and distributed through Microsoft Forms. It was pre-tested internally with people that had not seen the survey before to ensure clarity and relevance of the questions. Convenience sampling was used by distributing the survey through the SIB training mailing list, the SIB network, various international communities (among which ELIXIR (https://elixir-europe.org/) and Life Science Trainers (https://lifescitrainers.org/)) and social media. The survey was cross-sectional and open for responses for a period of 6 weeks, between March 18th and April 30th 2026, and participation was voluntary and anonymous. This manuscript follows the Consensus-Based Checklist for Reporting of Survey Studies (CROSS) [24]; the completed checklist is provided as a supplement.

### 2.2 Data preprocessing and missing value handling

All questions were mandatory, except for the Likert-type training needs questions. For these questions, respondents were explicitly permitted to skip topics that were not relevant to their work. Consequently, non-responses within the training needs questions were treated as meaningful omissions, i.e. ‘not relevant’. Therefore, missing values were imputed with the minimum ordinal score of 1, corresponding to a *“Not needed”* response. Three respondents who gave the same answers for all training-need topics were excluded from the clustering, because their similarity to other respondents is undefined. All data analysis steps were implemented in R ([25]; version 4.6.1) using the packages dplyr [26] for data manipulation and ggplot2 [27] for visualization. All the code can be found on GitHub (https://github.com/sib-swiss/analyze-training-needs-survey/) and Zenodo (https://zenodo.org/records/20446763).

### 2.3 Graph-based network and community detection

To define clusters of participants without relying on pre-existing demographic characteristics, we modeled the survey respondents using network analysis (Figure 1). In this network, each individual respondent is represented as a single node. For every possible pair of respondents, we calculated Spearman’s rank correlation coefficient based on their responses across all training sub-topics. We chose a rank-based correlation because Likert-type scores are ordinal, so the distances between answer categories cannot be assumed to be equal. This score reflects the similarity in their training needs. These similarity scores form the connections (edges) between nodes, where the weight of the connection represents the strength of the positive correlation between those two responses. To eliminate weak background noise and prevent the network from becoming too dense, we filtered based on a threshold of the Spearman correlation coefficient, pruning any connection falling below it. Unsupervised clustering of the filtered respondent network was then performed using the Leiden community detection algorithm [28]. The network analysis and community detection were conducted using the igraph package in R [29].

To find parameters with high cluster stability, we evaluated stability by resampling 80% of respondents over 100 iterations, reran the full network-construction and clustering procedure on each subsample, and compared the resulting partition to the full-data clustering (restricted to the subsampled respondents) using the Adjusted Rand Index (ARI), which ranges from 0 (chance-level agreement) to 1 (identical partitions). We swept this stability score over a grid of edge-threshold percentiles (50th–90th) and resolutions (0.5–2), restricting the comparison to configurations yielding a comparable number of interpretable clusters.

**Figure 1:**
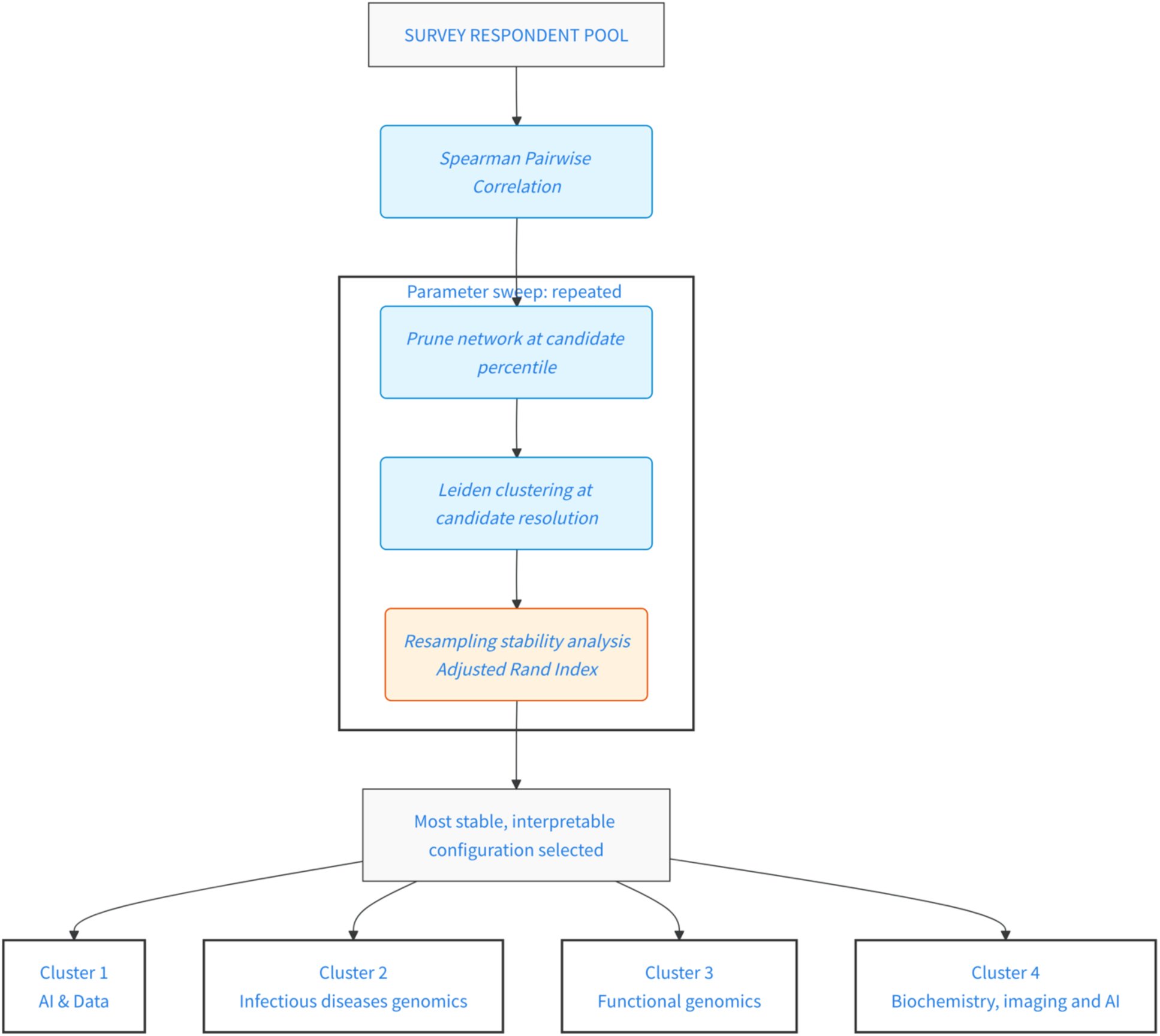
Graph-based clustering workflow for identifying learner profiles. Respondents are modeled as nodes in a network, with edges representing pairwise correlation of training needs. The edge-threshold percentile and resolution were identified using a parameter sweep. For each candidate combination, the network was pruned and clustered with Leiden community detection, and the resulting partition’s resampling stability (Adjusted Rand Index) was *evaluated. The most stable configuration from this sweep was then used to build the final pruned network and cluster respondents into the learner profiles reported here*.

## 3 Results

### 3.1 Survey data

A total of 260 responses were gathered, of which 176 (67.7%) were from Switzerland (Figure 2A). Most respondents from Switzerland were spread across six cantons that all have a large university (Figure 2 B). Most respondents were working in academia (n = 215, 82.7%; Figure 2 C). The career stage was distributed across academic professions (Figure 2 D), with a majority of PhD candidates (n = 77, 29.6%) and senior scientists/PIs (n = 67, 25.7%).

**Figure 2:**
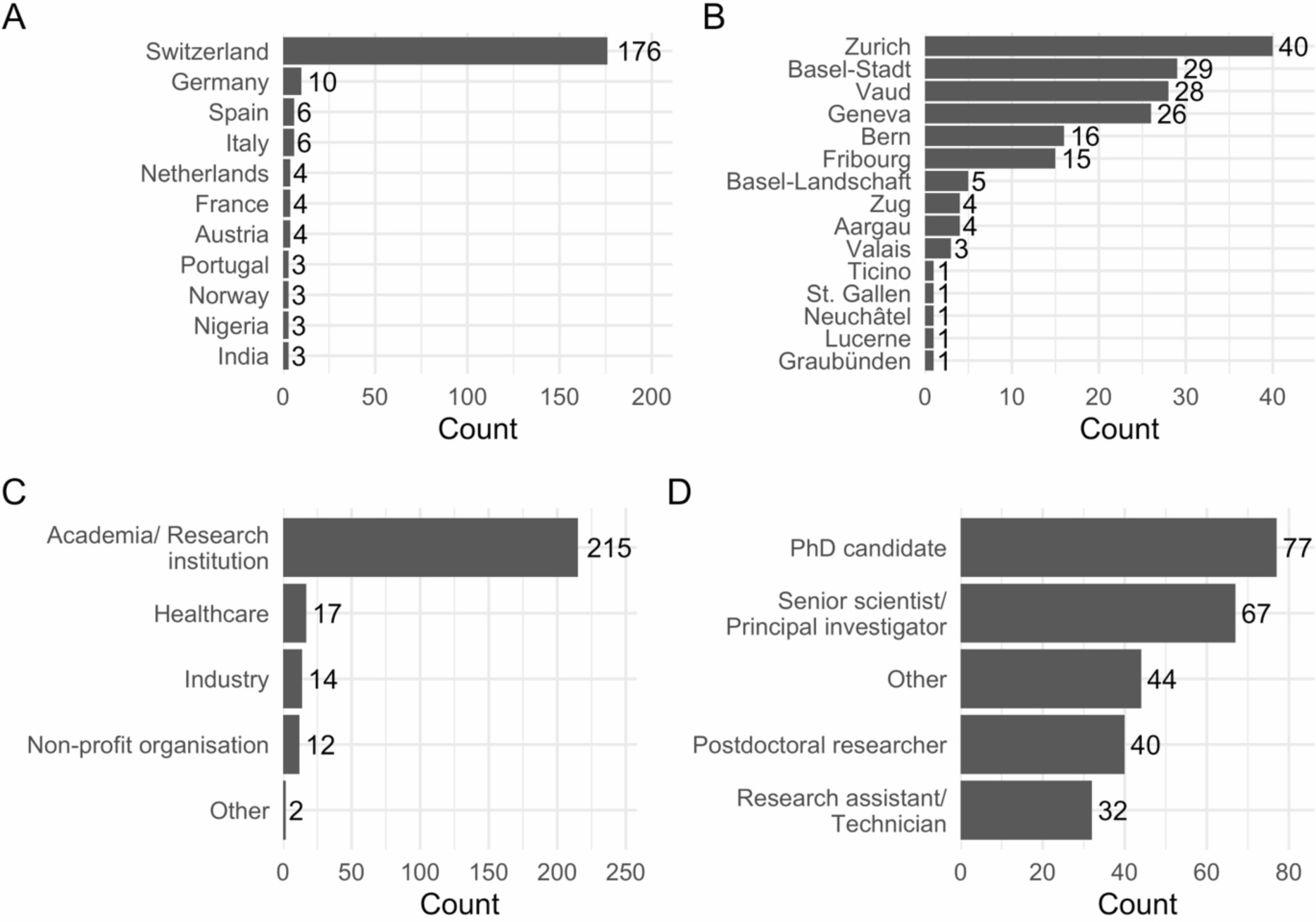
Respondent demographics. (A) Country of work. (B) Canton (if working in Switzerland). *(C) Organization type. (D) Career stage*.

Respondents were asked to indicate their training needs for 33 pre-defined course topics, grouped into five domains, on a 5-point Likert scale (Figure 3A). The topics most often rated “very needed” or “absolutely essential” were AI-related: AI-assisted coding and analysis (51%), using large language models for biomedical knowledge extraction (46%) and using AI for biomedicine (44%), together with making bioinformatics workflows findable and citable (45%) and sharing and reusing life science data (41%). At the other end, transposable elements analysis (14%), 16S sequencing data analysis (15%) and metabolic modelling (15%) were least often reported as needed.

Barriers to participation in SIB courses were also assessed using a 5-point Likert scale (Figure 3 B). For most of the listed factors, the majority of respondents reported no barrier or only a small one. The two most frequently reported barriers were distance to the course location and course fees, rated as a large or critical barrier by 33% and 25% of respondents respectively.

Respondents were also asked how much they liked each of nine delivery formats (Figure 3C; definitions in section 2.1). The most liked formats were blended learning (68% liked or strongly liked), multi-day intensive courses (63%), project-based learning (60%), short webinars (59%) and bring-your-own-data (56%). AI-assisted learning was the least liked (27%), and the only format that more respondents disliked (40%) than liked.

**Figure 3:**
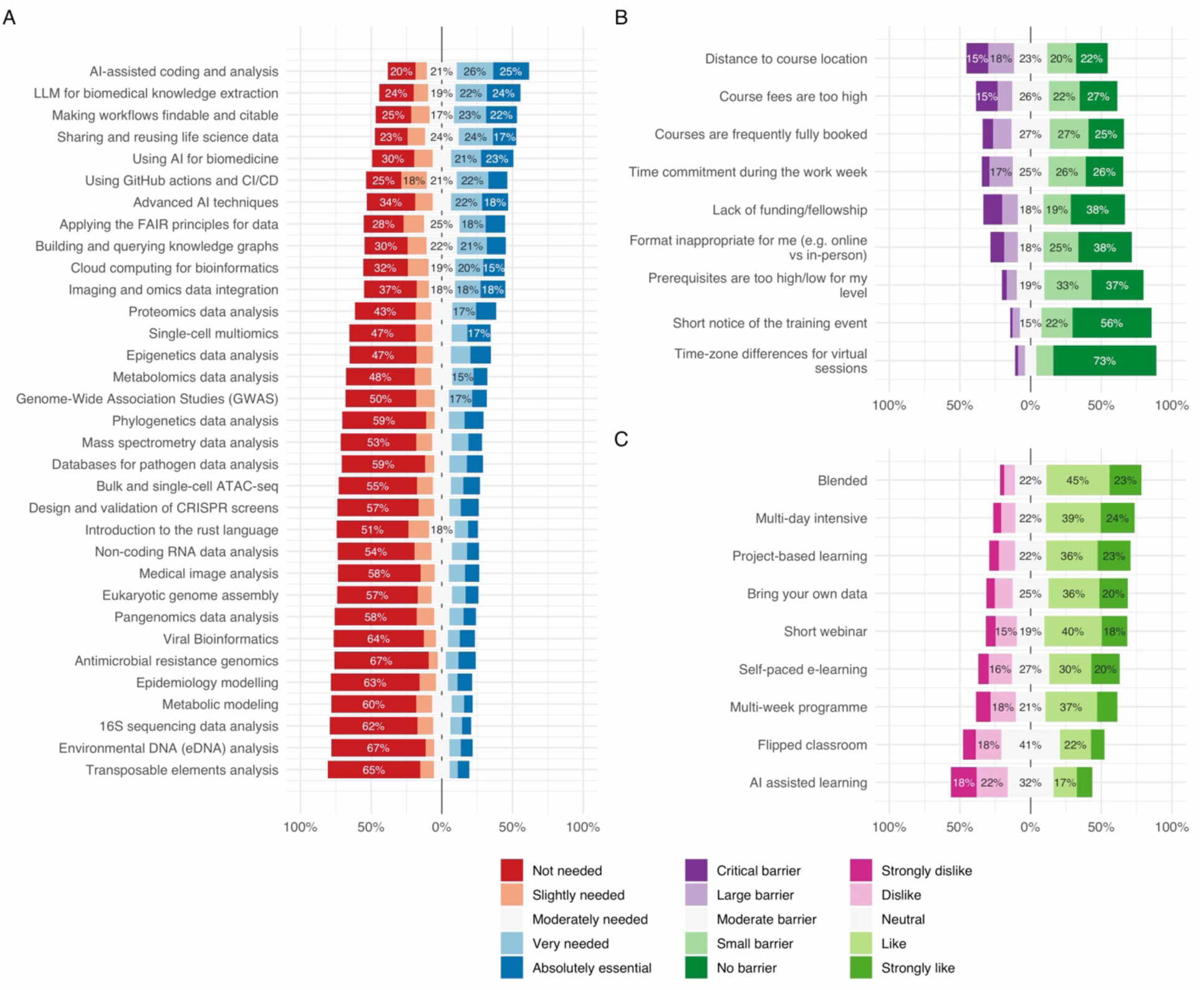
Overall responses of training needs, barriers and format preferences. (A) Training needs. (B) Barriers to participation in SIB courses. (C) Preferred learning formats. The topics, barriers and formats are ordered by the proportion of respondents who indicated that they needed the topic, that the factor was not a barrier, or that they liked the format, respectively. Skipped training-need topics were counted as ‘Not needed’ (see methods). Only percentages over 15% are shown. Some original answers were abbreviated for visualization purposes. For the full answers, please refer to Supplementary Data 1 & 2.

### 3.2 Graph-based clustering of course preference responses

Following the sweep of parameters (see section 2.3), the highest agreement was found for the combination 60th percentile) / resolution 1, resulting in a median Adjusted Rand Index (ARI) of 0.87, ranging from 0.50 to 0.91 (Figure 4). An ARI of 1 would indicate an identical partition on every resampling. At this threshold, connections were kept between respondents whose answers correlation above ρ = 0.181. This produced four clusters (n = 78, 50, 60 and 69). Recovery, which is the proportion of each cluster’s members that stayed together on resampling, was high for Clusters 1–3 (median recovery ≥ 0.98) and somewhat lower for Cluster 4 (median recovery 0.87; Table 1).

**Figure 4:**
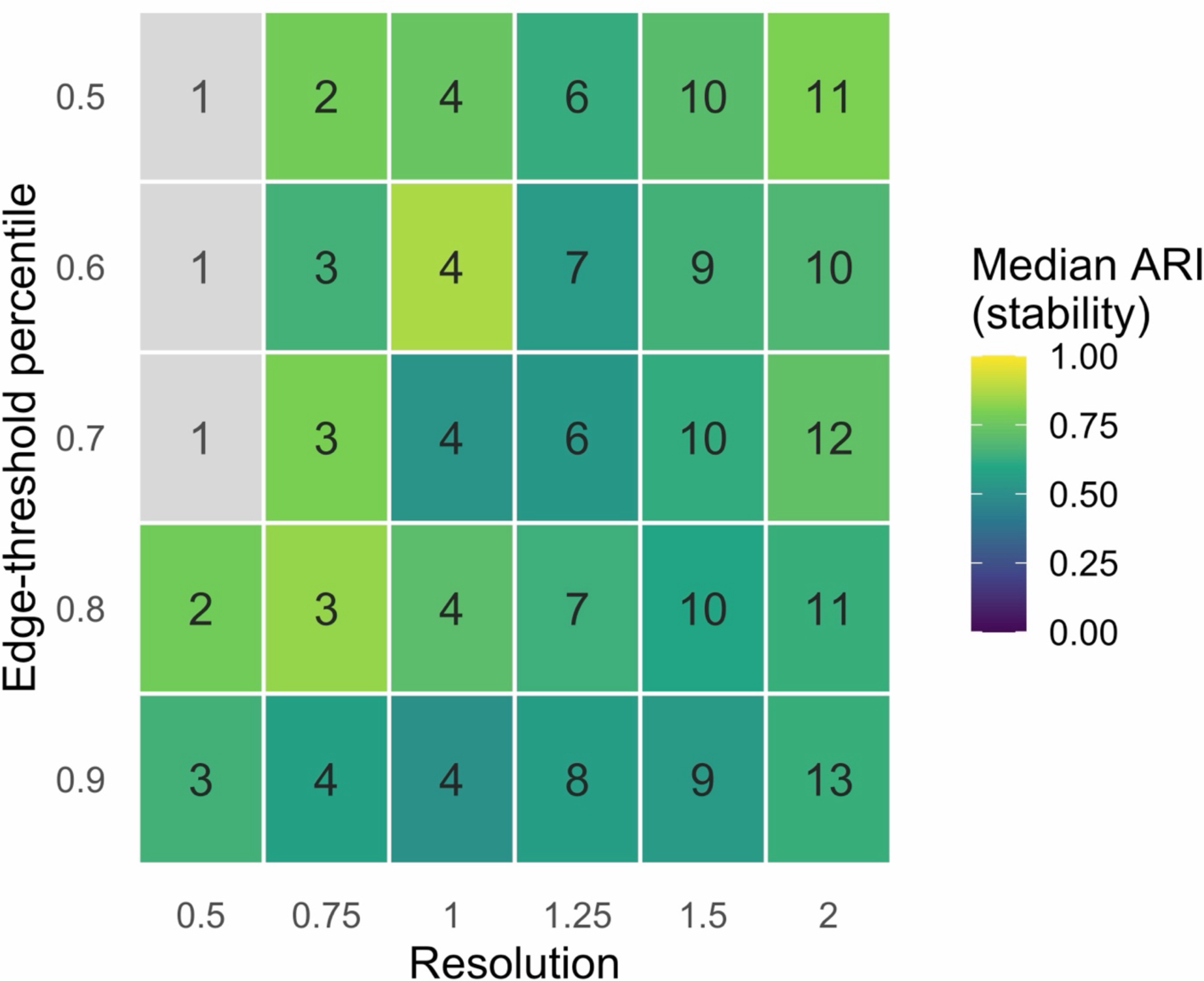
Resampling stability (median Adjusted Rand Index across 100 80%-subsample iterations) across a grid of edge-threshold percentiles and resolutions, using Leiden community detection. Numbers show the count of clusters at each setting; grey tiles indicate settings where the partition collapsed to a single cluster in some resampling iterations, making the Adjusted Rand Index undefined.

**Table 1:** Per-cluster recovery under resampling: for each learner-profile cluster, the fraction of its (sampled) members assigned to a single most-common cluster across 100 80%-subsample iterations.

| cluster | size<br>(respondents) | median<br>recovery | min<br>recovery | max<br>recovery |
|---|---|---|---|---|
| Cluster 1 | 78 | 0.991 | 0.881 | 1 |
| Cluster 2 | 50 | 0.977 | 0.953 | 0.977 |
| Cluster 3 | 60 | 0.989 | 0.938 | 1 |
| Cluster 4 | 69 | 0.868 | 0.608 | 0.891 |

### 3.3 Clusters and learner profiles

The four resulting clusters were characterized by distinct training needs, shown by their 10 most needed topics (Figure 5). This clearly shows that responses within each cluster have distinct preferences that are not visible when analyzing the entire respondent pool (Figure 3 A). Based on these distinct needs, we defined four learner profiles: Cluster 1 (Figure 5 A) is characterized by an interest in using AI and best practices for data, including application of LLMs and reproducible and FAIR data, and is hereafter referred to as the ‘AI and data’ profile. Cluster 2 (Figure 5 B) is characterized by infectious disease topics and possibly ecology, indicating an interest in phylogenetics, pathogen genomics and epidemiology, and is referred to as the ‘Infectious disease genomics’ profile. Cluster 3 (Figure 5 C) is characterized by an interest in functional genomics topics, indicating an interest in transcriptomics and epigenetics, and is referred to as the ‘Functional genomics’ profile. Cluster 4 (Figure 5 D) is characterized by training needs on biochemistry, imaging and AI, indicating an interest in proteomics, mass spectrometry and imaging and omics data integration, and is referred to as the ‘Biochemistry, imaging & AI’ profile. All clusters have topics related to AI in their top 10, indicating that AI is a cross-cutting theme across all learner profiles.

**Figure 5:**
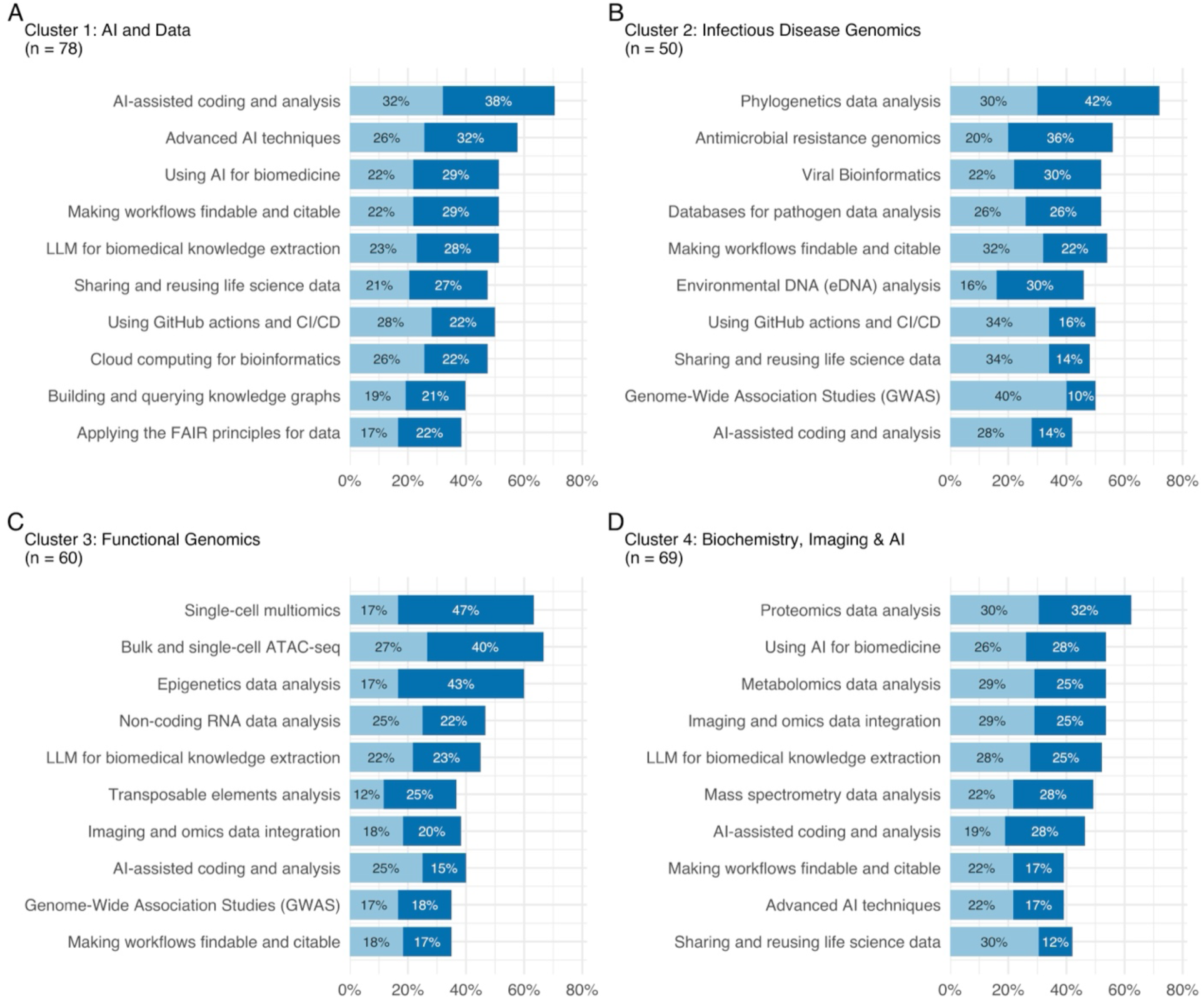
The top 10 most preferred topics are shown for each cluster, indicating that each cluster has distinct training needs. Only the answers “Very needed” (light blue) and “Absolutely essential” (dark blue) are shown. The numbers in the bars represent the count of respondents within the respective cluster selecting the answer. (A) Cluster 1: AI and data. (B) Cluster 2: Infectious disease genomics. (C) Cluster 3: Functional genomics. (D) Cluster 4: Biochemistry, imaging & AI. Some topics have been abbreviated for clarity; the full topic names presented to the respondents can be found in Supplementary Data 2.

The four profiles were also compared by demographic characteristics (career stage (Figure 6), sector, country and canton (Supplementary Figures 1-3)), barriers to participation (Supplementary Figure 4), and preferred learning formats (Supplementary Figure 5). The four profiles showed differences in career stage (Figure 6). Early-career scientists (PhD candidates and postdoctoral researchers) made up 51% of the ‘AI and data’ profile, compared with 48%, 45% and 34% in the other three, and senior scientists and principal investigators ranged from 18% in ‘AI and data’ to 30% in ‘Functional genomics’ and ‘Infectious Disease Genomics’. The clearest difference was in the ‘Infectious disease genomics’ profile, where 31% of respondents reported a career stage outside the listed categories, against 10–16% elsewhere. The share of respondents working in Switzerland ranged from 78% in ‘AI and data’ to 62% in ‘Functional genomics’ and ‘Biochemistry, Imaging and AI’ (Supplementary Figure 2). Barriers and format preferences also differed: course fees were most often reported as a large or critical barrier in the ‘Infectious disease genomics’ profile, and distance in the ‘Functional genomics’ profile (Supplementary Figure 4), while AI-assisted learning was the least liked format in all four profiles (Supplementary Figure 5).

**Figure 6:**
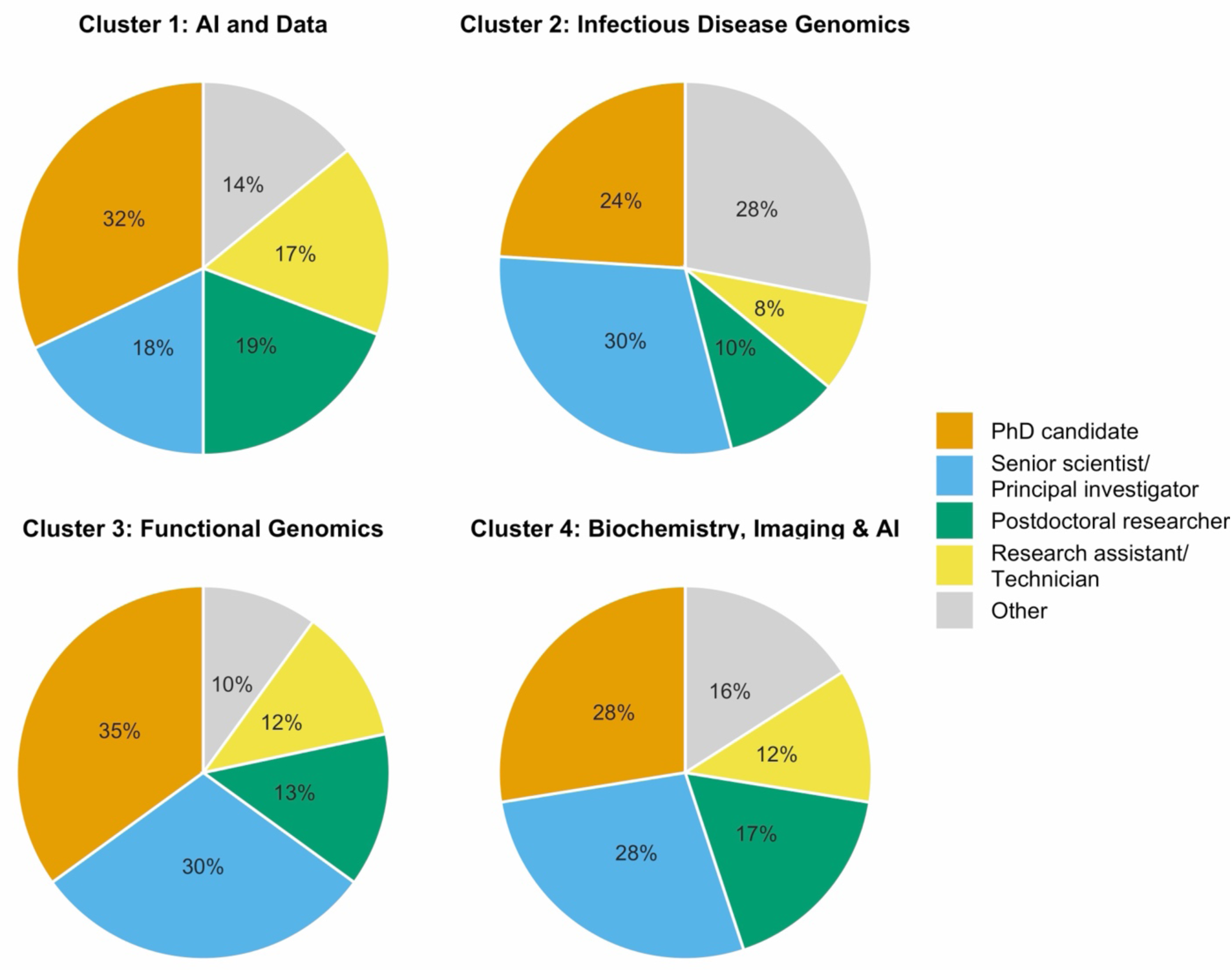
Career stage composition per cluster.

## 4 Discussion

This paper serves two purposes: it demonstrates a network-based clustering protocol for characterizing heterogeneous training needs from survey data, and it reports a snapshot of bioinformatics training needs in spring 2026, a period in which generative AI and large language models (LLMs) became increasingly relevant. In this section, we first discuss the methodological implications of our approach and then discuss the survey outcomes in the context of bioinformatics training.

### 4.1 Cluster-based analysis for survey data

The problems that can be solved with bioinformatics are very diverse, and professionals solving these problems are highly heterogeneous. Therefore, analyzing the survey as a homogeneous group, or slicing the group by demographic indicators only partly captures the diversity of training needs. Although examined here in the context of bioinformatics training, respondent clustering is a broadly applicable method for identifying meaningful patterns in survey data. For example, in citizenship norms [10], health literacy [11,12], bullying [14] or company training [13].

Several of these applications rely on latent class analysis [10,13,14] (LCA), a model-based alternative that assumes respondents are drawn from a mixture of latent groups, each characterized by its own probability of endorsing every item [30]. Unlike LCA, our approach does not require responses to different items to be conditionally independent once class membership is known. This assumption is hard to justify here, because people are often interested in several domains at once, and topics within the same domain tend to naturally go together. For example, in our data, all four learner profiles we identified had AI-related topics among their top 10 most-needed, a pattern consistent with a general interest in AI that cuts across profiles rather than being fully explained by profile membership, which is the kind of residual item correlation that the assumption of local independence rules out. Our approach also avoids the overfitting risk that LCA would carry here. As a rule of thumb, 300 responses or more are desirable for LCA to prevent overfitting [31], a number that can be quite difficult to reach for local surveys on training needs. In our case, we had four classes and 33 topics with five levels each. Estimating a full response-probability profile per item and class would require an estimated 531 free parameters (4 classes/clusters × 33 topics × 4 free category probabilities, plus 3 class-proportion parameters), exceeding our sample of 260 respondents and risking unstable or non-identifiable class solutions. Network-based clustering avoids this altogether by reducing each respondent’s profile to a single pairwise similarity score, so the number of parameters needed to define the clustering does not grow with the number of topics.

Our presented combination of survey with Likert-type questions and multiple-choice demographic questions with network-based clustering analysis provides an effective method to understand heterogeneous training needs. Similarly designed future training-needs surveys could run into similar limitations on the assumptions of LCA, making network-based clustering a broadly appropriate method for analysis. The clustering approach itself does not depend on the specific topics, wording, or number of domains used in our survey data — it only requires a set of Likert-type items sharing a common scale, asked of every respondent. This makes it directly reusable for other training-needs assessments. The accompanying implementation (https://github.com/sib-swiss/analyze-training-needs-survey) includes the survey design, preprocessing, and clustering steps as parameterized, reusable functions, so the protocol can be applied to a new dataset simply by supplying a new response export and question metadata file.

### 4.2 Survey outcomes

The majority of respondents were working in Switzerland as an academic researcher, likely because the survey was primarily advertised through the SIB training mailing list and other academic channels and networks within Switzerland, and participation was voluntary. This outcome aligns well with the aims of the survey, as SIB training is primarily intended for academic researchers working in Switzerland. Nevertheless, we think that the survey results can be used for similar training curricula in other countries. Here, we discuss the overall survey outcomes along the three dimensions of training needs, participation barriers and learning formats.

#### 4.2.1 A clear need for skills in AI and reproducible research

From the complete pool of survey results, it becomes clear that there is a demand for training on using (generative) AI and LLMs. Four of the topics were related to AI/LLM skills, and all four were in the top seven most needed topics. The two topics that were indicated as most needed were on how to use AI effectively: “AI-assisted coding and analysis”, and “Using large language models (LLM) for biomedical knowledge extraction”. A little bit less needed was the topic “Advanced AI techniques (agentic AI, MCP and RAGs)”, which is more related to developing AI tools. This shows that most respondents are aware of the potential use AI and LLMs in their daily work in bioinformatics, but there is a need for training on it. This perceived skill gap in AI and LLMs is a known barrier for the adoption and responsible use of AI in general [15,32], and it is reassuring to see that the respondents are aware of this gap and are interested in learning more about it, particularly in the ‘AI and data’ profile (Cluster 1).

The survey also revealed that there is a demand for training on FAIR data management and reproducible research, with “Making bioinformatics workflows findable and citable” and “Sharing and reusing life science data” as 3rd and 4th most needed respectively. This shows awareness of a need for learning about open and reproducible research practices amongst most respondents, which corresponds with earlier findings regarding perceptions on FAIR data practices [33], and its challenges [34].

While applying such outcomes to actual curriculum design, it is important to note that there can be a gap between intention - indicating a desire or need for training - and action - actually participating in a training course on the topic, known as the intention-behavior gap [35]. Although the questions on training needs were phrased in a way that the answer should relate to the respondents’ own training needs (“.. specify your training needs..”), some answers might represent what the respondents think their communities or colleagues need. This is also reflected in the relatively high proportion of senior scientists and PIs in the survey (25.7%), who might be more likely to answer on behalf of their group or lab rather than their own personal training needs. Therefore, the translation of these answers from intention to action, i.e. actually participating in a training course on the topic, might not be straightforward.

#### 4.2.2 Participation barriers

For most barriers, the majority of respondents did not perceive them as a barrier, or only a small one, to participation. Overall, the largest barriers were distance and course fees, reported as a large or critical barrier by 33% and 25% of respondents respectively, whereas barriers related to time-zone differences were perceived as minimal. The clustering protocol enabled the identification of limiting factors for the different learner profiles, rather than a single aggregate conclusion for the whole respondent pool, which is valuable to accommodate profile-specific barriers. Any resulting mitigation nevertheless has to weigh trade-offs the survey itself does not capture. Shifting more courses online, for instance, reduces a distance barrier but can come at some cost to student satisfaction [36] and learning effectiveness [37].

#### 4.2.3 The need for human interaction

Respondents particularly indicated a preference for short interactive live course formats, and combinations of both live and self-paced learning (blended). This shows the need of the human aspect in courses, such as the interaction between trainers and learners, and between learners themselves. Despite it being suggested to improve educational outcomes [38], there was a tendency towards dislike of AI-assisted learning formats, which is interesting given the high demand for training on AI and LLMs. This indicates that while respondents are interested in learning about AI, they prefer to learn in a more traditional format, with human interaction and guidance. This could be explained by overall trust issues regarding AI usage in education [39]. We do not know how respondents interpreted what ‘AI-assisted learning formats’ means. It is therefore important in next steps to understand how learners would appreciate this format, and what their expectations are.

#### 4.2.4 Learner profiles characterize heterogeneous training bioinformatics needs

By using graph-based clustering of the answers on training topics, we got a much more granular view, revealing insights that would otherwise remain hidden. For example, the topic “Using large language models (LLM) for biomedical knowledge extraction” was ranked second most needed by the complete pool of respondents. Within the profiles, though, it ranked well inside the top 10 for three of the four learner profiles, but 16th for the ‘Infectious disease genomics’ profile (Cluster 2). This shows that organizing a course on this topic would serve the primary needs of three learner profiles, but only partly those of Cluster 2’s respondents. The reverse also occurs. The topic “Phylogenetics data analysis” ranked 17th among the complete pool of respondents, suggesting a moderate to low priority. Within the ‘Infectious disease genomics’ profile (Cluster 2), however, it was the single most preferred topic (ranked first). Since the number of respondents was fairly evenly distributed across clusters, each learner profile deserves similar attention when designing a curriculum that matches these heterogeneous training needs. Organizing courses around profile-specific topics, rather than relying on the aggregate ranking alone, would therefore better serve substantial groups of respondents.

The learner profiles also enable a more targeted approach to address barriers to participation, learning formats and demographics. For example, a large proportion of respondents in the ‘Functional genomics’ profile (Cluster 3) indicated that distance to the course location is a barrier, and only a small proportion indicated that course fees are a barrier. On the contrary, a large proportion of respondents in the ‘Infectious disease genomics’ profile (Cluster 2) indicated that course fees are a barrier, and distance was the second most frequently indicated barrier. For curriculum design, this could suggest focussing on paid online multi-day courses for the ‘Functional genomics’ profile, and free webinars for the ‘Infectious disease genomics’ profile. This shows that these differences in barriers, preferred formats and demographics can inform the design of training programs that are tailored to the specific needs and preferences of each learner profile.

## 5 Conclusion

This study presents a network-based clustering protocol for characterizing heterogeneous training needs using Likert-type survey questions in combination with multiple-choice demographic questions and demonstrates its use on the 2026 SIB training-needs survey. By clustering respondents based on their shared needs rather than demographics, we identified four meaningful learner profiles (AI and data, infectious disease genomics, functional genomics, and biochemistry, imaging and AI), each with distinct topic priorities, participation barriers, and format preferences. For curriculum design this argues for building courses around learner profiles rather than around a single ranked list of topics. Across all profiles, the analysis indicated a clear need for training on AI and reproducible data skills alongside a preference for human-led, interactive formats over AI-assisted learning, raising questions about trust and the role of human interaction in AI-era training. As the protocol depends only on a shared Likert scale across survey items, we expect it to generalize to other training-needs assessments. The full pipeline is openly available to support such reuse, alongside the parameter-sensitivity and algorithm-choice considerations discussed above.

