## Supplementary Figures for "Characterization of heterogeneous bioinformatics training needs in the AI era"

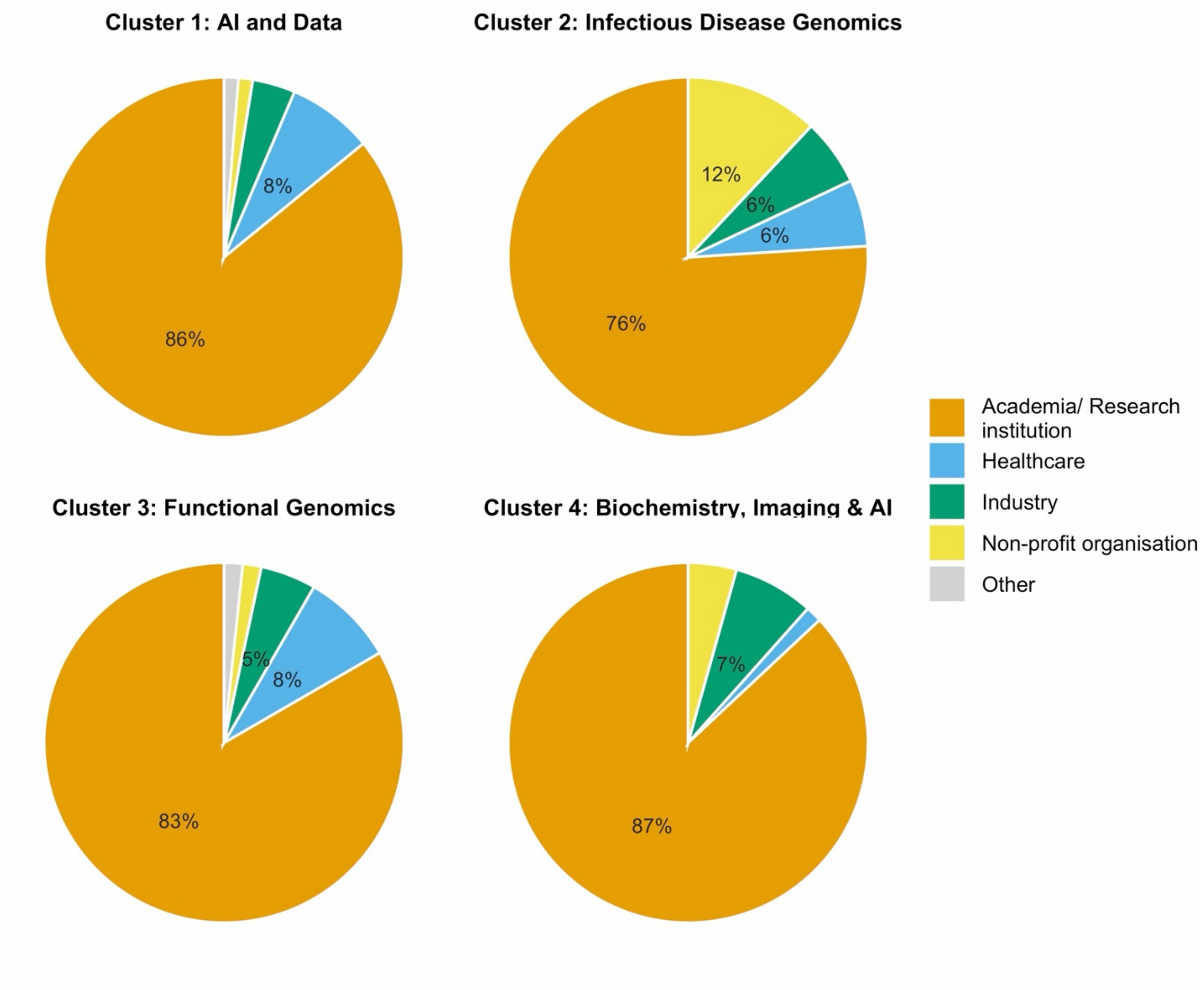


Supplementary Figure 1. Profile composition per sector

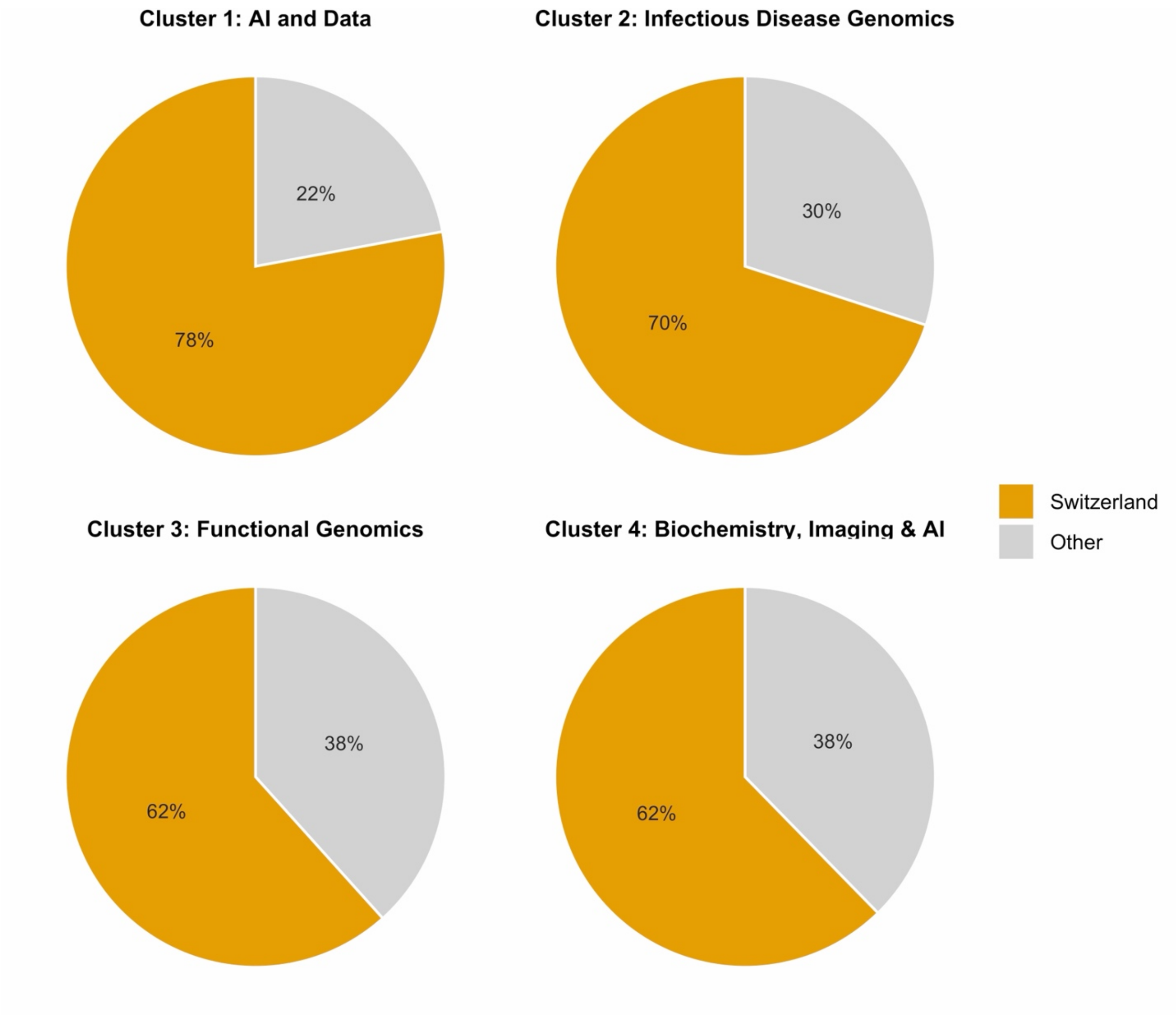


Supplementary Figure 2. Profile composition Switzerland versus other countries.

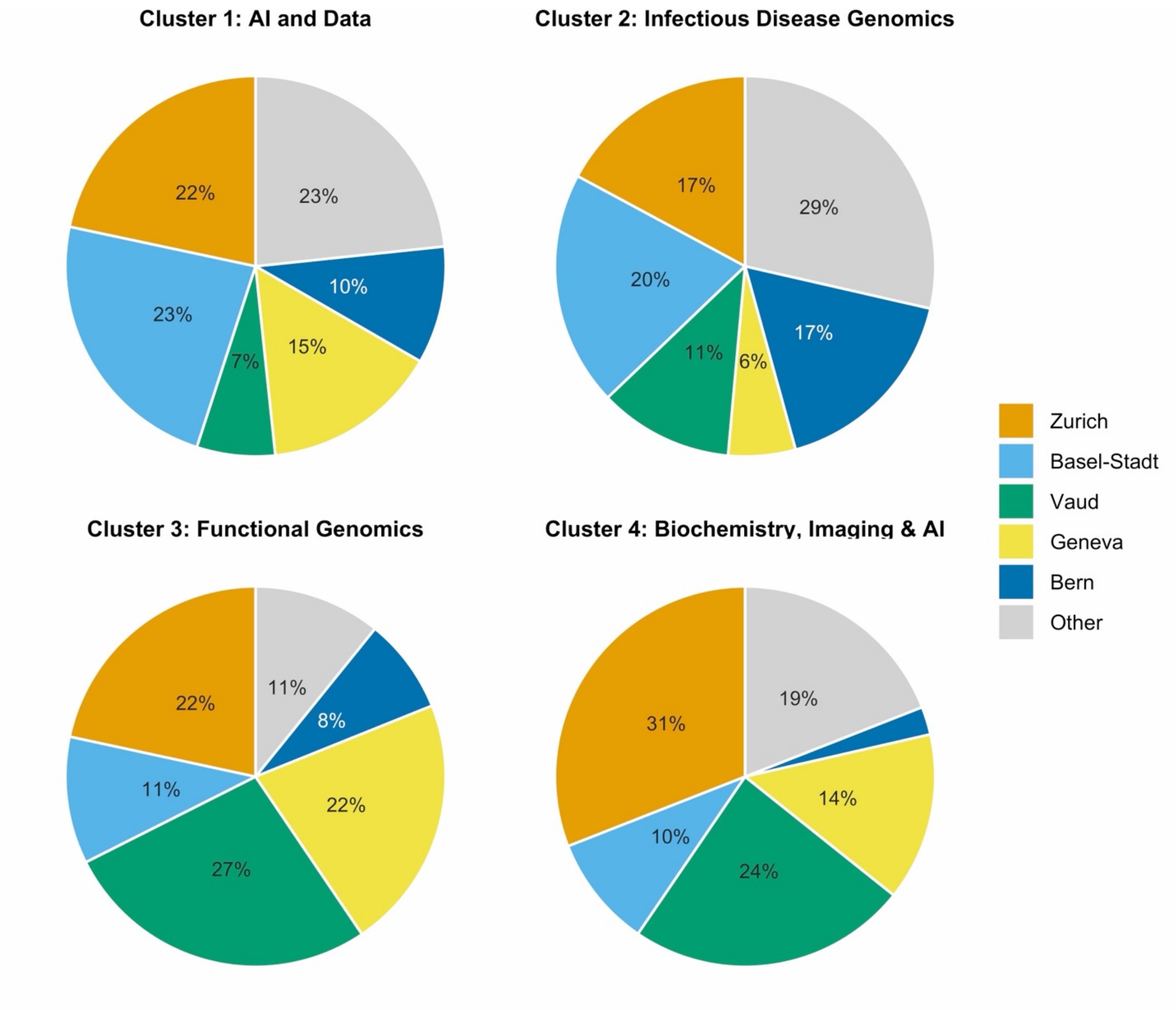


Supplementary Figure 3. Profile composition per canton. Only respondents who indicated to work in Switzerland are represented.

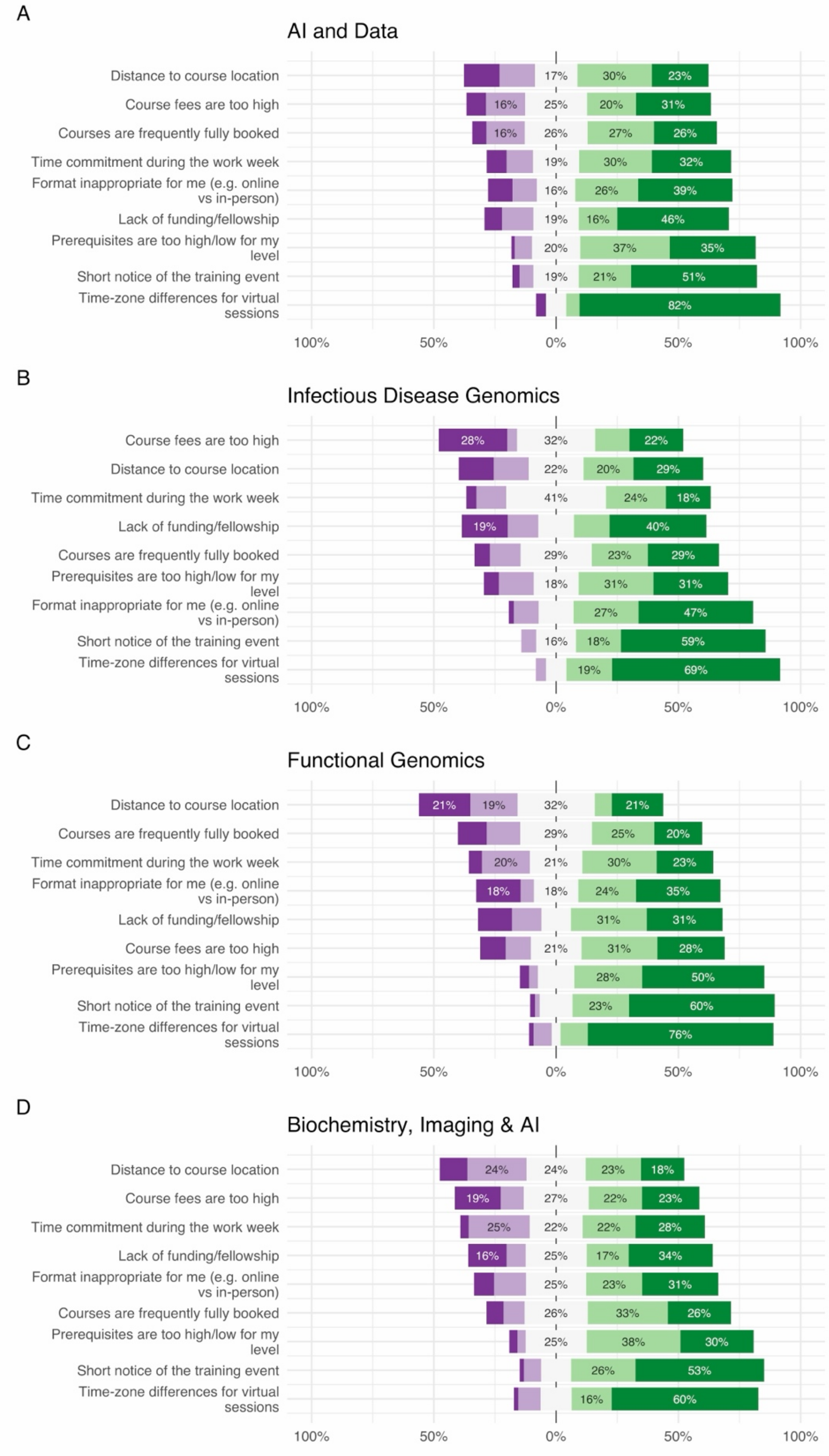


Supplementary Figure 4. Barriers per profile. Colors as in figure 3.

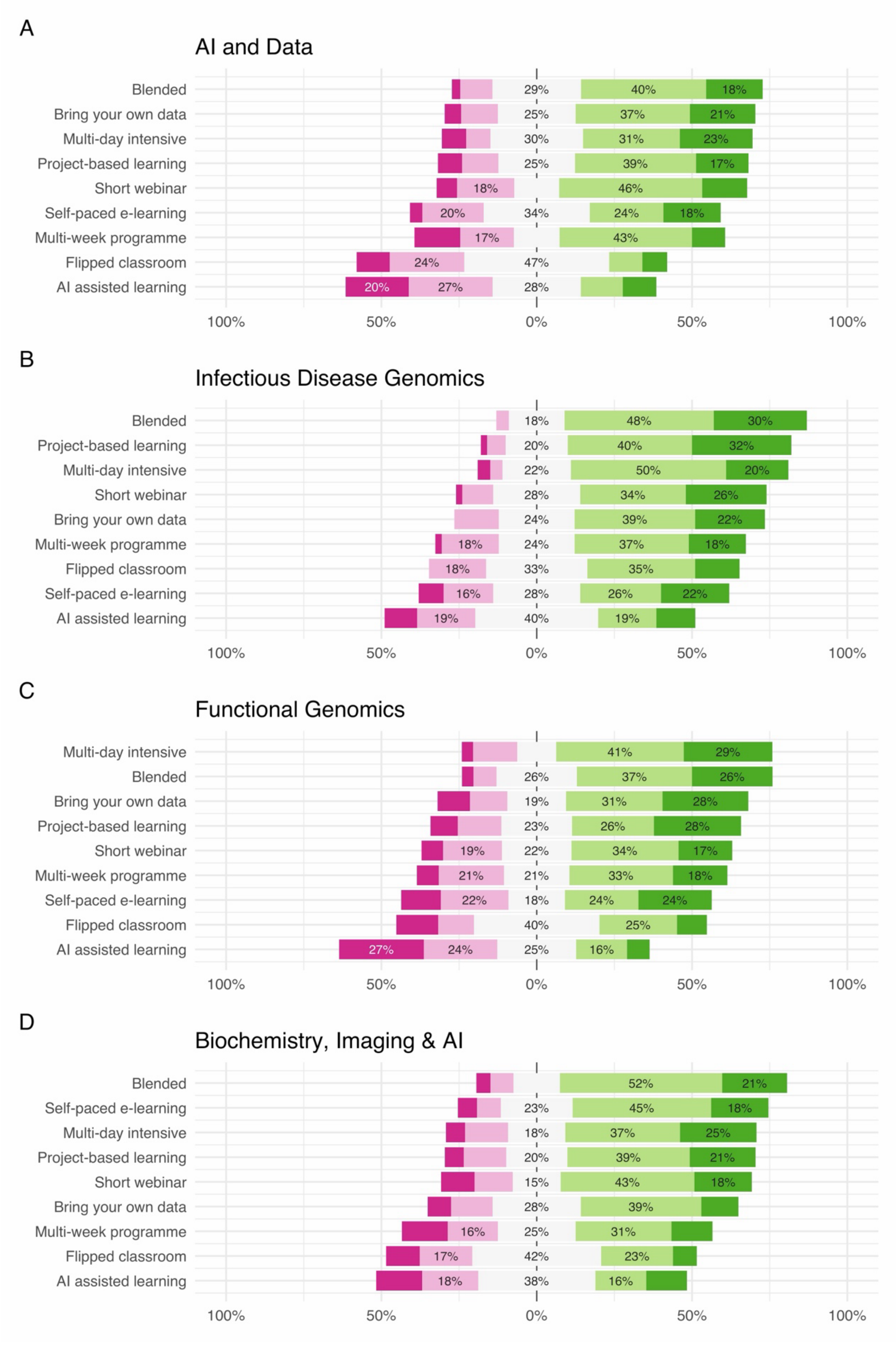


Supplementary Figure 5. Preferred learning formats per profile. Colors as in figure 3.
